# Effects of the social environment on movement-integrated habitat selection

**DOI:** 10.1101/2021.02.11.430740

**Authors:** Quinn M.R. Webber, Christina M. Prokopenko, Katrien A. Kingdon, Julie W. Turner, Eric Vander Wal

## Abstract

1. Movement links the distribution of habitats with the social environment of animals using those habitats; yet integrating movement, habitat selection, and socioecology remains an opportunity for further study.
2. Here, our objective was to disentangle the roles of habitat selection and social association as drivers of collective movement in a gregarious ungulate. To accomplish this objective, we (1) assessed whether socially familiar individuals form discrete social communities and whether social communities have high spatial, but not necessarily temporal, overlap; and (2) we modelled the relationship between collective movement and selection of foraging habitats using socially informed integrated step selection analysis.
3. We used social network analysis to assign individuals to social communities and determine short and long-term social preference among individuals. Using integrated step selection functions (iSSF), we then modelled the effect of social processes, i.e., nearest neighbour distance and social preference, and movement behaviour on patterns of habitat selection.
4. Based on assignment of individuals to social communities and home range overlap analyses, individuals assorted into discrete social communities, and these communities had high spatial overlap. By unifying social network analysis with iSSF, we identified movement-dependent social association, where individuals foraged with more familiar individuals, but moved collectively with any between foraging patches.
5. Our study demonstrates that social behaviour and space use are inter-related based on spatial overlap of social communities and movement-dependent habitat selection. Movement, habitat selection, and social behaviour are linked in theory. Here, we put these concepts into practice to demonstrate that movement is the glue connecting individual habitat selection to the social environment.

## 1. Introduction

Movement is defined by a change in spatial location and is the behavioural link between the physical space an animal occupies and the resources available to them (Van Moorter, Rolandsen, Basille, &Gaillard, 2016). In the context of the social environment, movement represents the connection between the distribution of resources and the social structure of animals that consume those resources (He, Maldonado-Chaparro, &Farine, 2019). Disentangling the social and spatial drivers of movement is a formidable challenge within behavioural ecology. In many cases, research omits the social contexts within which animals move to, from, and within the areas that contain foraging resources (Spiegel, Leu, Bull, &Sih, 2017; Strandburg-Peshkin, Papageorgiou, Crofoot, &Farine, 2018). Spatially-explicit models of sociality highlight that some gregarious species aggregate at areas associated with profitable foraging resources (Chamaillé-Jammes, Fritz, Valeix, Murindagomo, &Clobert, 2008), whereas some territorial species only interact at territory edges (Spiegel, Sih, Leu, &Bull, 2018). Sharing space, either at foraging sites, territory edges, or elsewhere within an animal’s range is required to form the social environment. For example, animals are predicted to select habitat as a function of the profitability and availability of the habitat (van Beest et al., 2014). A logical extension can be made to conspecifics; individuals form groups based on their familiarity with conspecifics and the profitability of associating with familiar conspecifics. We aim to quantify the relative importance of habitat and conspecifics by developing a socially informed integrated step selection analysis, a movement-based method that accounts for the relative intensity of selection for habitats and neighbours.

For social animals, individual movement shapes social encounters and subsequent interactions with conspecifics that can affect collective movement (Jolles, King, &Killen, 2020). Further complicating our understanding of collective movement is the idea that the type, quality, and distribution of habitats on the landscape can constrain or promote collective movement (Strandburg-Peshkin, Farine, Crofoot, &Couzin, 2017). For example, dense vegetation impedes visibility, which could reduce the probability a group remains together. In addition, individual movement and habitat selection are affected by the distribution of resources. For example, patchily distributed foraging resources could facilitate large aggregations, whereas homogenously distributed foraging resources could result in a reduction in social associations (Spiegel, Leu, Bull, &Sih, 2017). The physical space an individual, or group, occupies and the distribution and availability of foraging resources within that space are important drivers of animal movement and the social environment an individual experiences (He et al., 2019).

Animals typically select for habitats that maximize foraging and minimize risk of predation; an important trade-off because most habitats do not accommodate both high quality foraging *and* low predation risk. When animals aggregate in large groups, the per capita risk of predation is lower. Thus, animals in larger groups reduce time spent vigilant (Creel, Schuette, &Christianson, 2014). Furthermore, individuals in larger groups tend to select more risky habitats, including foraging in open areas (Lima, 1995). However, not all social groups are equal; some groups contain unfamiliar individuals (i.e., anonymous groups) (Harel, Spiegel, Getz, &Nathan, 2017), while others contain familiar individuals (Lachlan, Crooks, &Laland, 1998). For anonymous and familiar groups, social foraging occurs when the costs and benefits of an individual’s foraging behaviour are linked with the foraging behaviour of conspecifics (Giraldeau &Dubois, 2008). Social foraging can be most beneficial when social information about foraging resources comes from familiar individuals (Patin, Fortin, Sueur, &Chamaillé-Jammes, 2019). For example, when foraging resources are unpredictable, familiar individuals obtain reliable information from conspecifics to increase foraging efficiency (Jones, Patrick, Evans, &Wells, 2020; Spiegel &Crofoot, 2016), such that time searching for forage is reduced in favour of more time spent foraging. In the context of movement and habitat selection, theory on social foraging and the benefits of social familiarity provides a framework through which the costs and benefits of collective movement can be explored (Giraldeau &Caraco, 2018; Giraldeau &Dubois, 2008).

Apparent social familiarity or preference is the long-term repeated social association due to shared space at the same time. Although individuals often interact with many conspecifics, non-random repeated social interactions or associations with certain individuals form the basis for social preference (Mourier, Vercelloni, &Planes, 2012). Proximately, long-term social relationships can influence collective movement via the reliability of information transfer about foraging resources or predator risk (Best, Seddon, Dwyer, &Goldizen, 2013; Muller, Cantor, Cuthill, &Harris, 2018), while ultimately they can enhance fitness (Silk, 2007). The social environment can be influenced by the availability of foraging resources, but social communities can also be composed of individuals with similar physiological or nutritional requirements that occupy the same locations. Apparent social preference may therefore arise as a function of spatial constraints (Spiegel, Leu, Sih, &Bull, 2016), including physical barriers, such as rivers or mountains. Disentangling social preference from spatial constraint could inform our understanding of collective movement and habitat selection (Croft, Darden, &Wey, 2016; Pinter-Wollman et al., 2013).

Here, we develop a unified framework to bridge the gap between social network analysis and movement ecology. We disentangle the roles of social preference and collective movement on habitat selection behaviour by parameterizing socially informed integrated step selection models (Figure 1). Animal social networks often comprise distinct sub-networks, or social communities, defined by the existence of social preference among discrete clusters of individuals (Mourier et al., 2012). Using a social ungulate as a model system, our objective was to disentangle the roles of habitat selection and social association as drivers of collective movement in a gregarious ungulate (*Rangifer tarandus*) when the availability and distribution of foraging resources are variable. We calculated three distinct measures of social preference. First, we assigned individuals to social communities based on a community detection algorithm. Second, we assessed the temporal stability of social association among individuals. Third, we estimated spatial overlap of social communities using home range analyses. Due to variance in the distribution of foraging resources on the landscape, we expected that access to social information via close proximity to conspecifics should influence patterns of selection for foraging resources. Specifically, individuals with stronger social preference should select foraging habitat collectively. The corollary is that individuals should also take short steps in the presence of conspecifics, given that from a movement ecology perspective, shorter steps typically represent foraging behaviour and longer steps represent searching behaviour (Owen-Smith, Fryxell, &Merrill, 2010).

**Figure 1.**
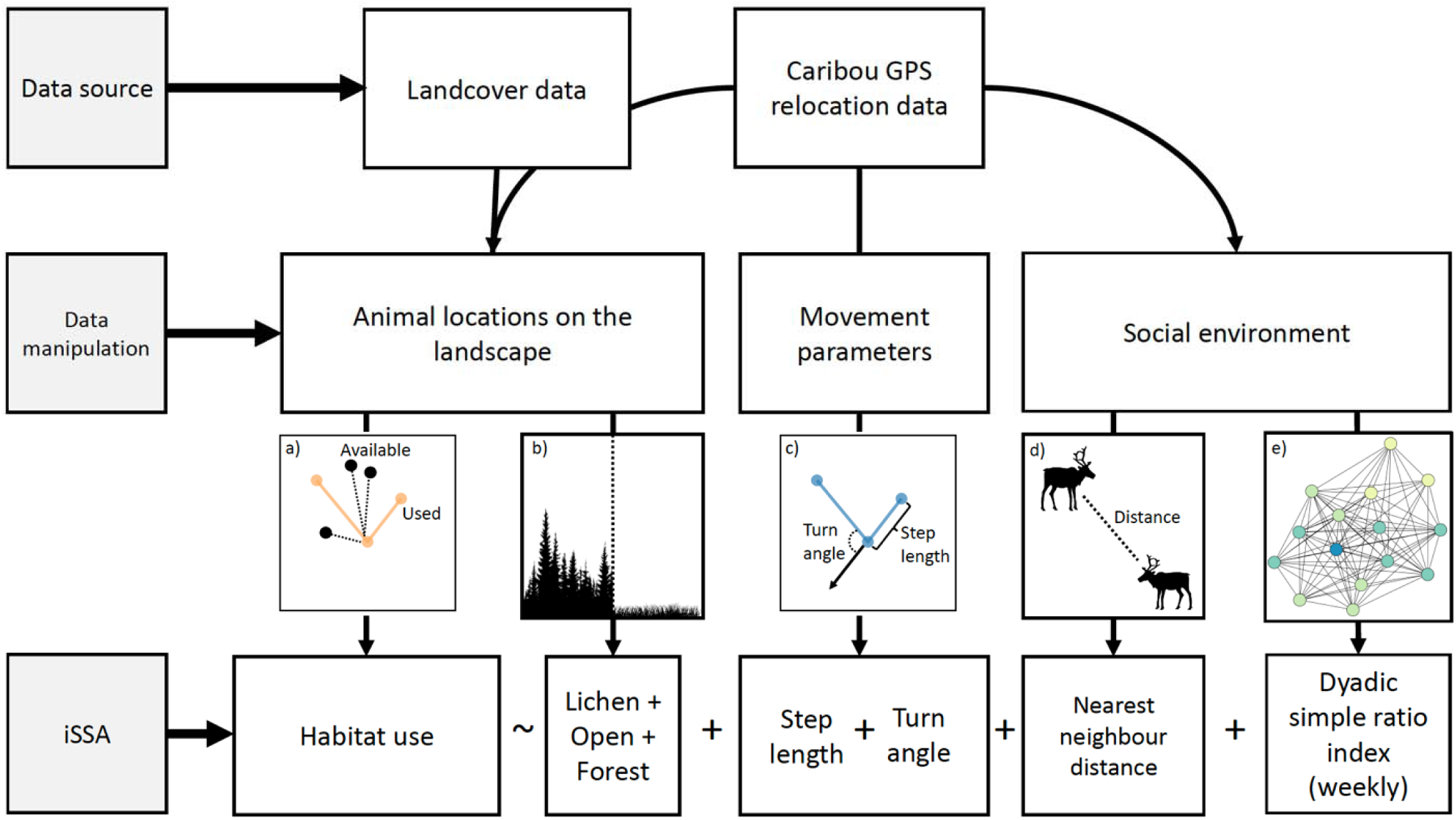
Summary of the data pipeline used to generate integrated step selection function (iSSF) models. Primary data sources were landcover data and caribou GPS relocation data, which were combined to determine the physical locations of animals on the landscape. The pairing of animal locations and landcover data was used to generate the comparison of used to available points (panel a), which is the response variable in iSSF models, as well as the habitat type in which a given relocation occurred: lichen (defined in text as open-forage), open (defined in text as open-movement), and forest (panel b). Caribou relocation data were also used to generate two movement parameters (panel c) and aspects of the social environment (panels d and f). Movement parameters included turn angle, which is the angular deviation between the headings of two consecutive steps, and step length, which is the linear distance between consecutive relocations. The social environment included nearest neighbour distance (panel d) and weekly social networks and the dyadic simple ratio index generated based on a moving-window as a proxy for short-term social preference (panel e). The bottom row represents a graphical formulation of our iSSF models, where habitat selection (1:10 ratio of used to available relocations) was regressed against habitat type (lichen, open, and forest), movement parameters (step length and turn angle), nearest neighbour distance, and weekly dyadic simple ratio index.

## 2. Materials and Methods

### 2.1 Caribou as a model system

We investigated patterns of movement, space use, and social behaviour for caribou (*Rangifer tarandus*) on Fogo Island, Newfoundland, Canada. Fogo Island is a small (∼237km^2^) island off the northeastern coast of Newfoundland with a humid continental climate (see Supplementary Materials for details). Between 1964-1967, 26 caribou were introduced to Fogo Island from the Island of Newfoundland (Bergerud &Mercer, 1989). Currently, Fogo Island has a population of approximately 300 caribou (Newfoundland and Labrador Wildlife Division, unpublished data). Caribou live in fission-fusion societies (Lesmerises, Johnson, &St-Laurent, 2018), and throughout much of their range, caribou forage primarily on lichen, grasses, sedges, and other deciduous browse with access to these resources changing between the seasons (Bergerud, 1974). During winter (January to March), the landscape is covered by snow, and caribou forage primarily on lichen (Webber, Ferraro, Hendrix, &Vander Wal, 2022). Lichen is heterogeneously distributed, and access is impeded by snow and ice cover. Caribou dig holes in the snow, termed craters, to access lichen in the winter, often where snow depth is relatively shallow (∼30–60 cm deep). Consequently, caribou have limited access to lichen buried under the snow and tend to re-use established craters. To cope with this limitation, caribou use conspecific attraction and social information transfer to gain access to foraging opportunities (Peignier et al., 2019). In addition, caribou typically avoid forested habitats due to deep snow in forests and lack of access to forage opportunities (Fortin, Courtois, Etcheverry, Dussault, &Gingras, 2008), whereas most open habitats on Fogo Island are windswept in the winter, facilitating foraging and movement (Bergerud, 1974).

We used GPS location data collected from Fogo Island caribou (2017–2019) to assess the relationship between social behaviour, habitat selection, and movement (see supplementary information for details on collaring procedures). For all analyses, we restricted locations to only include relocations from the first 75 days of each year (1 January–16 March). Each relocation was assigned to a given habitat classification that was extracted from Landsat images with 30m × 30m pixels (Integrated-Informatics, 2014). Locations were categorized as one of open foraging (lichen barrens), open moving (wetland, rocky outcrops, and water/ice), or forest (conifer scrub, mixed wood, and conifer forest). We then calculated the proportion of each habitat type (i.e., open foraging, open moving, or forest) within 200 m around each used and available point location (see below). Adult female caribou (n = 26 individual caribou, n = 72 caribou-years) were immobilized and fitted with global positioning system (GPS) collars (Lotek Wireless Inc., Newmarket, ON, Canada, GPS4400M collars, 1,250 g). Prior to analyses, we removed all erroneous and outlier GPS locations following Bjørneraas et al. (Bjørneraas, Van Moorter, Rolandsen, &Herfindal, 2010). We did not collar all female caribou in the herds, however, and collared individuals were randomly selected from the population. We therefore assume that our sample of collared animals was randomly distributed. Although associations between collared and uncollared animals were unrecorded, we assumed that our networks (see below) were unbiased representations of the relative degree of social association among all caribou. All animal captures and handling procedures were consistent with the American Society of Mammologist guidelines and were approved by Memorial University Animal Use Protocol No. 20152067.

### 2.2 Formulating integrated step selection models

Integrated step selection function (iSSF) simultaneously incorporates movement and habitat selection within a conditional logistic regression framework (Figure 1) (Avgar, Potts, Lewis, &Boyce, 2016; Basille et al., 2015; Duchesne, Fortin, &Rivest, 2015). As in other resource and step selection analyses (Fortin et al., 2005), iSSF models habitat selection as a binomial response variable where ‘use’ represents the location an animal was observed and ‘availability’ represents the geographical area an animal could potentially use but was not necessarily observed (Figure S1). iSSF defines availability based on empirically fitted distributions of step lengths and turn angles (Avgar et al., 2016), where a step is the linear connection between consecutive relocations, and turn angle is the angular deviation between the headings of two consecutive steps (Prokopenko, Boyce, &Avgar, 2017). We generated available steps and turn angles based on the distributions informed by observed population-level movement behaviour using the *amt* package in R (Signer, Fieberg, &Avgar, 2019). First, we sampled step lengths from a gamma distribution of observed step lengths for the study population; values were log-transformed for analysis. The statistical coefficient of log-transformed step length is a modifier of the shape parameter from the gamma distribution originally used to generate available steps (Avgar et al., 2016). Second, we sampled turn angles (measured in radians) for available steps from observed values between -rr and rr following a Von Mises distribution. Each observed relocation was paired through a shared start point with 20 available steps generated from step-length and turn-angle distributions and compared in a conditional logistic regression framework (see section 2.7). In addition to generating available movement parameters, we also generated an available social environment (see below). To evaluate the predictive performance of our model, we used k-fold (k = 5) cross validation (Roberts et al., 2017) following the methods of Fortin et al. (2009). For details on k-fold cross validation see Appendix 2.

### 2.3 Social network analysis

We used the R (R Core Team, 2019) packages *spatsoc* (Robitaille, Webber, &Vander Wal, 2019) and *igraph* (Csárdi &Nepusz, 2006) to generate proximity-based social association networks from GPS location data. Nodes in the networks represented individual caribou and edges represented the frequency of association based on proximity between individuals. We generated social networks at two scales based on proximity of locations between individual caribou: (1) seasonal winter networks to assign individuals to social communities and assess long-term social preference and (2) weekly networks to assess the role of short-term social preference on patterns of habitat selection (section 2.2). Social communities represent a subset of individuals within a network that are more closely connected with each other than with the rest of the network. For networks at both seasonal and weekly scales, we assumed association between two individuals when simultaneous locations (i.e. GPS relocations that occurred within 5 minutes of each other) were within 50 m of one another (Lesmerises et al., 2018; Peignier et al., 2019). We selected the 50 m threshold based on the standard distance applied to assign individuals to groups in studies of ungulate group size and social behaviour (Kasozi &Montgomery, 2020). We applied the ‘chain rule’, where each discrete GPS fix was buffered by 50 m and we considered individuals in the same group if 50 m buffers for two or more individuals were contiguous, even if some individuals were beyond 50 m of one another. We weighted edges of social networks by the strength of association between dyads of caribou using the simple ratio index (Cairns &Schwager, 1987), SRI:

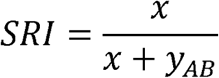

where *x* is the number of times individuals A and B were within 50 m of each other and *y*_*AB*_ is the number of simultaneous fixes from individuals A and B that were separated by >50 m (Farine &Whitehead, 2015).

### 2.4 Detecting social communities: long-term social preference

For seasonal winter social networks, we used a community detection algorithm to define social communities (Newman, 2006). We assessed social community structure for each winter to determine the broadest extent of social structure. Modularity is a commonly used measure that defines how well-connected social communities are to one another. It is calculated from the weighted proportion of edges that occur within a community, minus the expected proportion of edges, if edges were distributed randomly in the network (Newman, 2006). A modularity value close to 1 indicates a network with a strong clustered structure in which interactions of individuals belonging to different clusters do not occur. We quantified modularity (*Q*) for observed annual winter networks. To ensure observed social structure did not occur at random, we compared these values to null models (Spiegel et al., 2016). Specifically, we generated null models based on GPS fixes to reduce potential for type II error typically associated with node-based permutations (Farine, 2014). Following Spiegel et al. (2016), we re-ordered daily GPS movement trajectories for each individual while maintaining the temporal path sequence within each time block (e.g., day 1 and day 2 may be swapped). This technique is a robust network randomization procedure for GPS data because: 1) it maintains the spatial aspects of an individual’s movement; 2) by randomizing movement trajectories of individuals independent of one another, temporal dependencies of movement are decoupled (Spiegel et al., 2016). We repeated this procedure 100 times for annual winter networks and re-calculated modularity at each iteration. We then compared observed modularity (*Q*) values to the null distribution and determined whether the observed *Q* value fell within the 95% confidence interval of the distribution of *Q* values (Mourier et al., 2012).

In addition to comparing observed *Q* values from annual winter networks to a null distribution, we also calculated a community assortativity coefficient (R_com_) to assess confidence in the assignment of an individual to a given community (Shizuka &Farine, 2016). Specifically, R_com_ = 0 indicates no confidence in the assignment of an individual to a community, while R_com_= 1 indicates certainty in the assignment of an individual to its community.

### 2.5 Weekly networks and lagged association rates: short-term social preference

We iteratively generated weekly social networks using a moving window approach and calculated the observed SRI to be included as a covariate in our iSSF model (see section 2.2). The first network was calculated for 1 January to 7 January, the second was 2 January to 8 January, and so on. Weekly networks contained 84 relocations per individual (12 relocations per day). For each of these networks, we used dyadic values of SRI as a proxy for short-term social preference. We used a three-step process. First, to incorporate SRI within the iSSF framework, we determined the identity and distance (m) of each individual’s nearest neighbour at each relocation. Second, for each focal individual and their nearest neighbour at each relocation, we matched the dyadic SRI value for the prior week. For example, for individual A at 12:00 on 8 January, we determined the nearest neighbour was individual B and we extracted the dyadic SRI value for these individuals for the previous week. Third, we repeated steps one and two for all ‘available’ relocations defined by random steps generated in the iSSF (section 2.2). Therefore, each individual at each relocation had an observed weekly dyadic SRI value and a series of available weekly dyadic SRI values (see section 2.2).

In addition to incorporating social preference directly within the iSSF model, we also assessed social preference by estimating within-season temporal patterns in associations between individuals by calculating the lagged association rate (LAR). We calculated the LAR for social networks using the *asnipe* package in R (Farine, 2013). LARs measure the probability that pairs of individuals associating at a given relocation would still associate at subsequent relocations (Whitehead, 2008). We generated annual LARs to compare temporal stability to assess potential for within-season patterns of association among individuals. In addition, we also compared seasonal LARs for individuals in the same annual winter social community to LARs for individuals in different annual winter social communities to assess potential for within-season patterns of association among individuals (Figure S4).

### 2.6 Home range overlap between social communities

To determine spatial overlap of social communities we estimated home ranges for winter social communities using the area of the 95% isopleths from fixed kernel density estimates (Worton, 1989) for each social community in each year with the *href* smoothing parameter in the *adehabitatHR* package in R. Data from all individuals in a given social community were pooled to estimate the community home range. We estimated home range overlap between social communities with the utilization distribution overlap index (UDOI), where higher values of UDOI represent a greater proportion of overlap and lower values represent lower proportion of overlap (Fieberg &Kochanny, 2005).

### 2.7 Modelling collective movement and habitat selection

We fit a single iSSF model with a series of fixed and random effects using the *glmmTMB* package in R following Muff et al. (Muff, Signer, &Fieberg, 2020). We took advantage of the fact that the conditional logistic regression model is a likelihood-equivalent to a Poisson model with stratum□specific fixed intercepts. The approach outlined by Muff et al. (2020) uses a mixed modelling approach which allows intercepts and/or slopes to vary by individual, while also incorporating shared information that is present in the data from different individuals (Fieberg, Rieger, Zicus, &Schildcrout, 2009). For social species that may move collectively, and therefore have correlated movement trajectories, varying intercepts by individual is recommended to account for correlation within nested groupings of locations (Hebblewhite &Merrill, 2008). Following Muff et al. (2020), all variables included in the fixed effect structure were also included in the random effect structure. Our model included the proportion of lichen, forest, and open habitat within 200 m of the point location, the natural log-transformed step length, natural log-transformed nearest neighbour distance, and weekly dyadic simple ratio index (section 2.3). Nearest neighbour distance (m) was measured as the distance between a focal individual and the nearest collared conspecific and was calculated for all used and available steps. We also included interactions between step length and each of the proportion of lichen, forest, and open habitats within 200 m of the point location, nearest neighbour distance and step length, and simple ratio index, nearest neighbour distance and each of the proportion of lichen, forest, and open habitats within 200 m of the point location, and simple ratio index and each of the proportion of lichen, forest, and open habitats within 200 m of the point location (see Table S1). For interactions that included nearest neighbour distance, we used either distance at the start of a step or at the end of the step, depending on the other variable in the interaction (Figure S1). Specifically, for the interaction between step length and nearest neighbour distance, we used distance at the start of the step because the likelihood of taking a shorter or longer step is predicted to vary based on the distance to conspecifics before the step is taken. By contrast, for interactions between habitat variables and nearest neighbour distance, we used distance at the end of the step because the likelihood of selecting a given habitat is predicted to vary based on the distance to conspecifics when that habitat is being selected, i.e., at the end of the step.

### 2.8 Calculating effect sizes

We calculated individual-level relative selection strength (RSS) to demonstrate how habitat features influenced selection (Avgar, Lele, Keim, &Boyce, 2017). We calculated the strength for selecting one step over another that differed in the habitat value where those steps ended. RSS was calculated for each habitat type (i.e., forest, lichen, or open habitats) as a function of nearest neighbour distance and the shared dyadic simple ratio index between nearest neighbours.

## 3. Results

We found that individuals associated with members of multiple communities, and associations were stronger among members of a given community. Depending on the year, social networks comprised 2–6 social communities, and although community assortativity (R_com_) was similar across years, there was high certainty (range = 0.95–1.00) of an individual’s assignment to a given community in a given year (Table S1). In addition, lagged association rates (LAR) within each winter confirmed temporal stability of community assortment, where association rates for members of the same winter community remained higher than association rates for members of different communities in each year (Figure 2). Seasonal winter values of modularity (*Q*) were significantly lower than the distribution of *Q* generated from null models (Figure S2), suggesting that social networks were structured weakly into communities with frequent inter-community social associations (Table S1). In support of our expectation, we observed relatively high spatial overlap between different winter social communities (average UDOI = 0.37, SD = 0.34, range = 0–0.98; Figure S3; Table S2), thus facilitating the potential for association between social communities.

**Figure 2.**
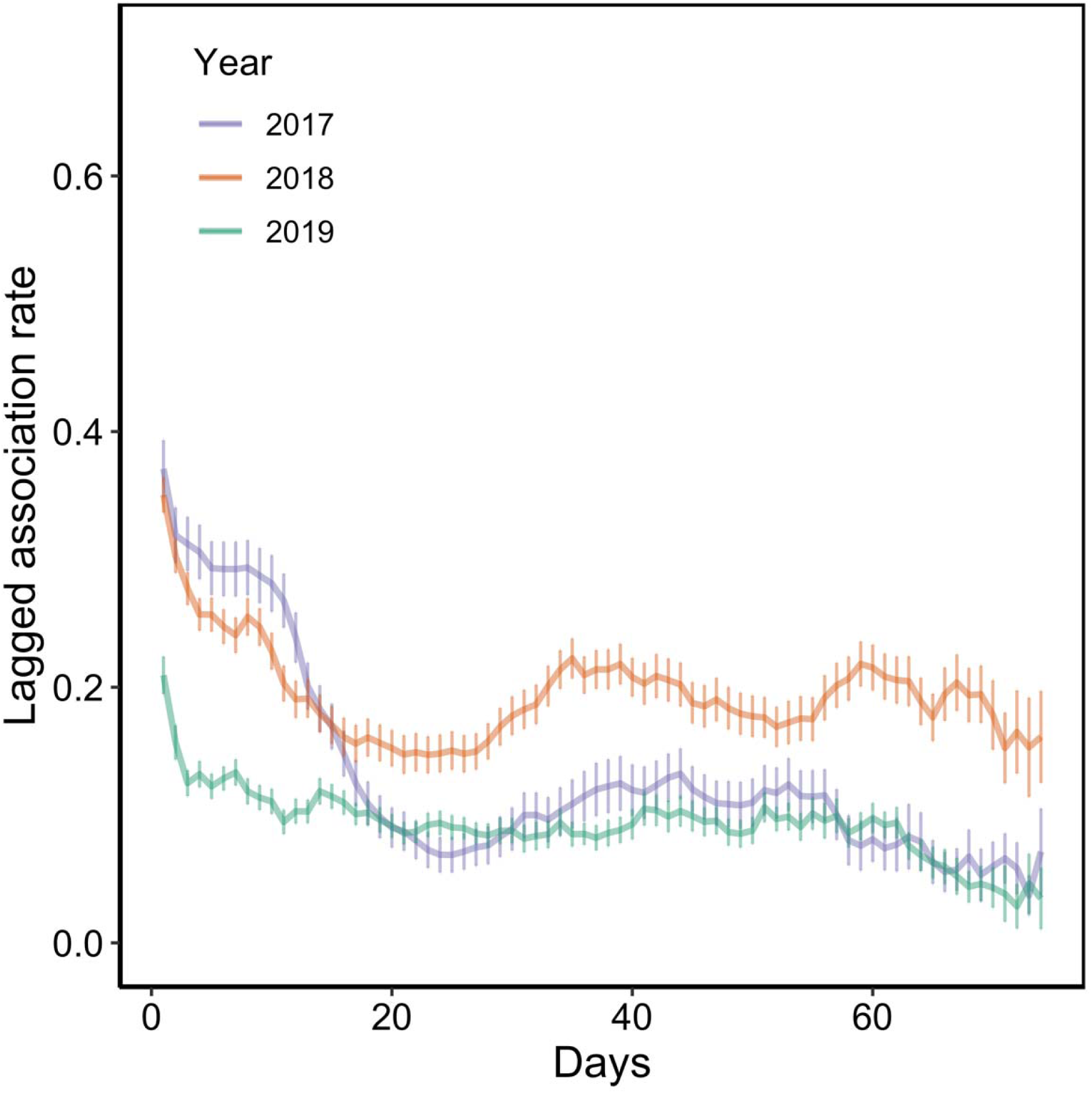
Observed annual lagged association rate (LAR) for caribou, calculated as the probability that any pair of individuals associated on a given day, are still associated on subsequent days. Note, the time period for LAR analysis was 1 January to 16 March. Error bars represent the standard error of all pairwise association rates calculated on each day.

Overall, we found that caribou are highly social in nearly all circumstances and that caribou prefer to select all habitats with familiar conspecifics (Figure 4). Despite these findings, the effect of the social environment on selection was nuanced, and we found partial support for our expectation of social foraging. Individuals moved more slowly when selecting lichen and when they shared a high SRI value with their nearest neighbour, suggesting potential that conspecific familiarity influenced foraging-related movement (Table S2). However, relative to its availability, caribou moved more quickly through open habitat, perhaps to travel between foraging sites (Figure 3). Meanwhile, relative selection strength for all habitats decayed as nearest neighbours were further away, however, relative selection for lichen habitat was stronger than forest and open habitats (Figure 4). Our k-fold cross-validation had high scores (rho = 0.80 SE ± 0.06), demonstrating our model was better than random at predicting where caribou moved (see Figure S6 for coefficients for variables in each fold).

**Figure 3.**
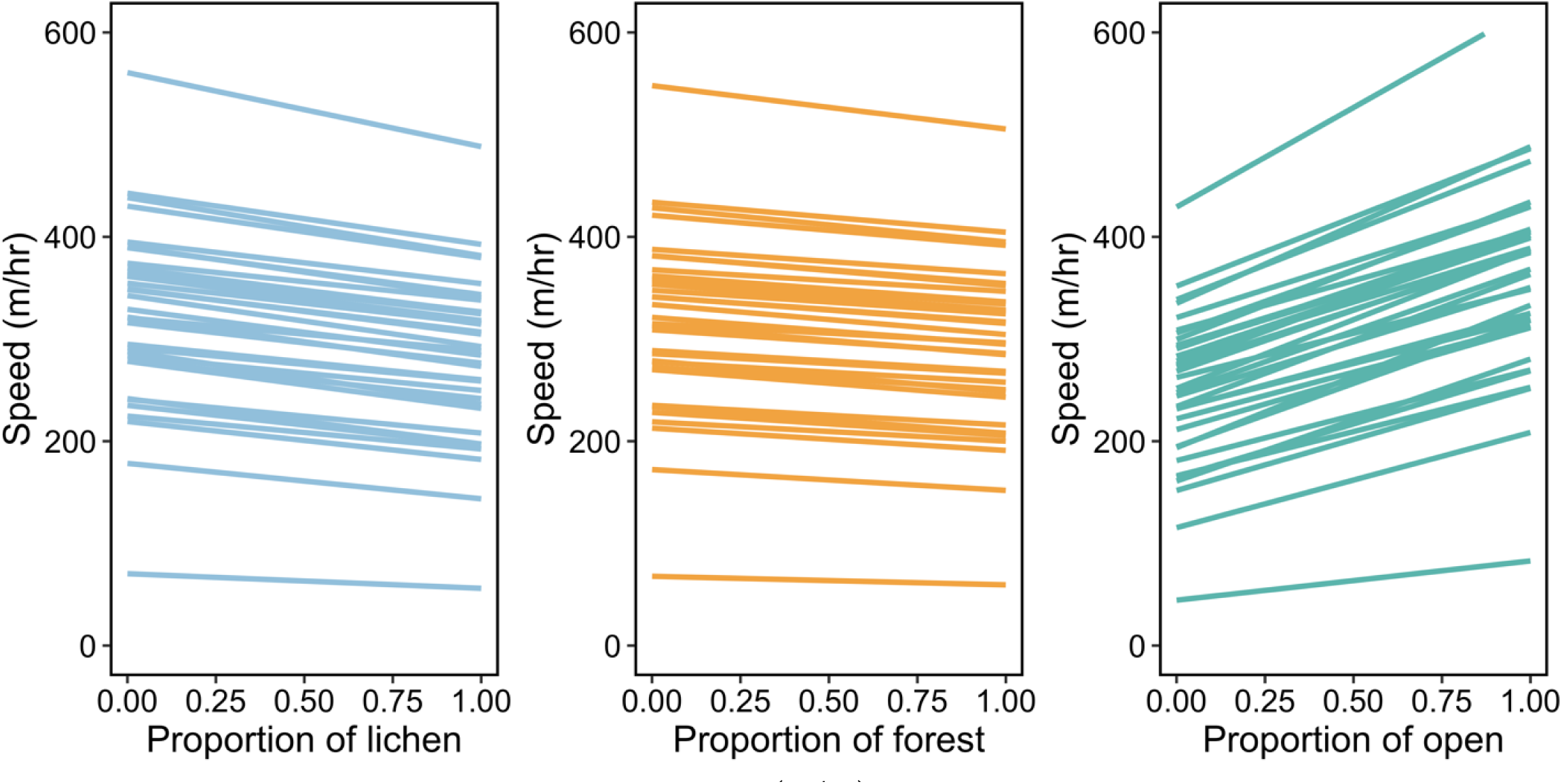
Relationship between expected speed (m/hr) as a function of changes in the proportion of lichen, forest, and open habitats within 200 m of a given point location for individual caribou.

**Figure 4.**
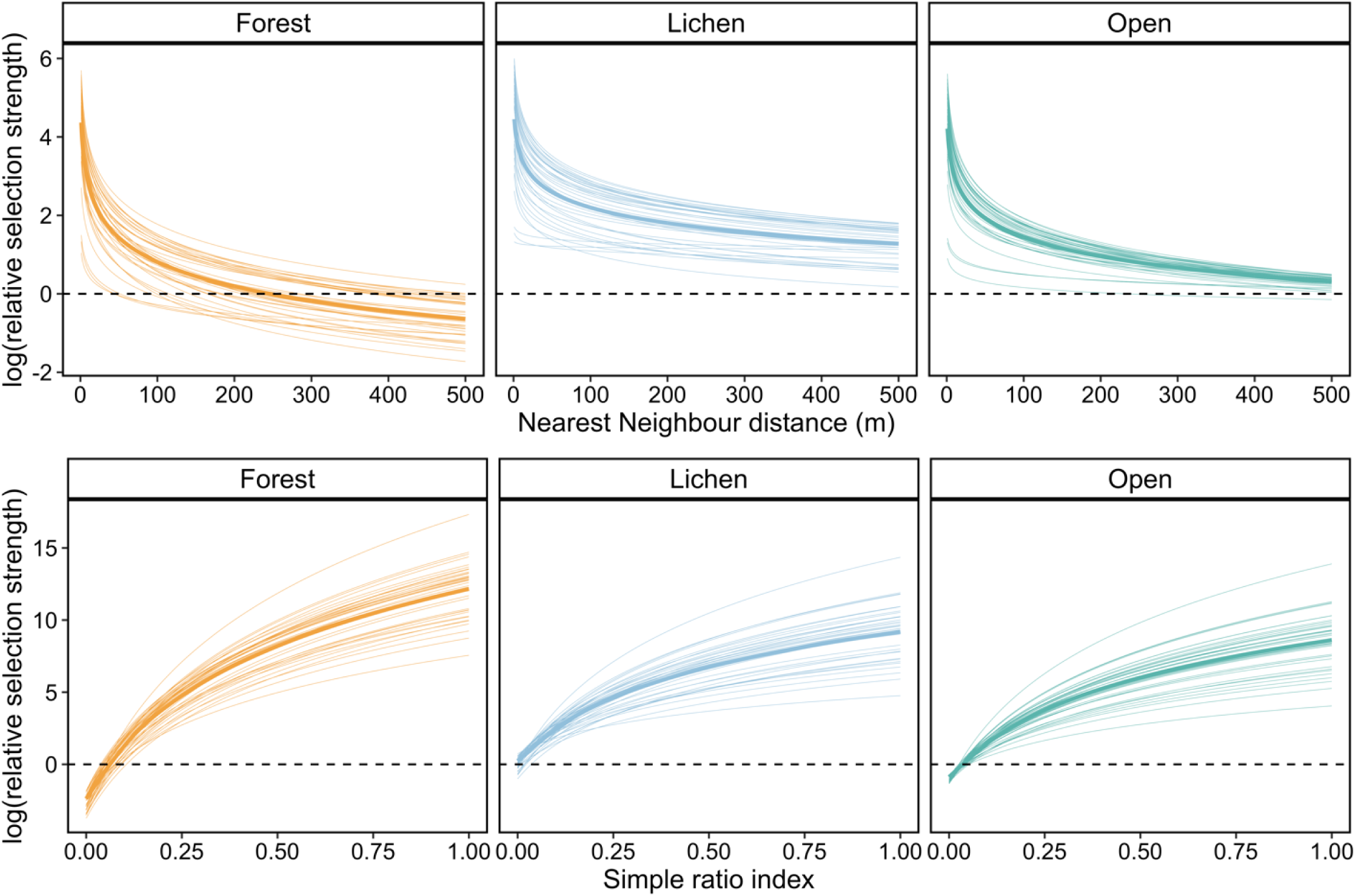
Relative selection strength of forest, lichen, and open habitats as a function of nearest neighbour distance (m) (top panels) and shared dyadic simple ration index between nearest neighbours (bottom panels). The dotted horizontal line represents no response, while values above the line indicate the population is selecting to be closer to that habitat than expected or to have higher shared dyadic simple ratio index than expected, and below the dotted line the population is selecting to be farther from that habitat than expected or to have a lower shared dyadic simple ratio index than expected. Interpretation for RSS values are that individuals generally tend to select to be near to conspecifics when selecting lichen and open habitats relative to their availability, whereas the response to nearest neighbours in forest habitat relative to its availability is limited. Meanwhile, individuals select for familiar nearest neighbours in all habitats.

## 4. Discussion

Our study examined apparent social preference in the context of shared space use using socially informed integrated step selection functions. We present a framework that unifies social networks within a traditional movement ecology and habitat selection framework. Although individual social associations were well mixed at the population level, we found that social networks were structured into discrete communities. Despite spatial overlap between different social communities, which suggests an opportunity for individuals to interact with members of other communities, we highlight two forms of within-community social preference, including long-term temporal stability of associations among individuals, and an effect of short-term social preference on habitat selection. Further, we found that individuals tended to select foraging habitat near familiar individuals but moved between foraging habitats with conspecifics regardless of their degree of familiarity, suggesting the social environment can vary relative to the speed animals are moving. The processes underlying community structure appear to be social, and not spatial. Based on our unification of social network analysis with integrated step selection functions, we highlight the influence of collective movement and preferred associations on habitat selection and foraging.

Testing social preference as a driver of movement and habitat selection required establishing the existence of discrete communities and long-term social associations within the population-level network. Indeed, the formation of social communities, in combination with our lagged association analysis, confirmed the existence of temporal stability in social associations for members of the same social community. The loose formation of non-random social communities is consistent with expectations of fission-fusion dynamics, where groups merge and split through space and time (Sueur et al., 2011). Community formation was driven in part by social preference, but aspects of space use, including shared space, could also influence the formation of social communities, even if they are relatively weak (Daizaburo Shizuka et al., 2014). We found high spatial overlap between social communities, suggesting that physical barriers on the landscape do not explain the formation of discrete social communities. For social communities to emerge from a well-mixed population, individuals in different communities must have high spatial, but low temporal overlap in shared geographical space, thus revealing the importance of space and time in the formation of social communities (Cantor et al., 2012). Disentangling space and time within the social environment reveals distinct social communities and groups of individuals that are more likely to associate than by chance (Spiegel et al., 2016). On resource limited landscapes, individuals are expected to aggregate in close proximity to those resources, for example, elephants (*Loxodonta africana*) aggregate near water-holes, which are a limiting resource (Chamaillé-Jammes et al., 2008). At the population-level, social networks were highly connected, thus providing the impetus to quantify socially informed patterns in movement and habitat selection.

Our findings reveal that caribou are social in nearly all circumstances, although we observed a social hierarchy of movement-dependent social associations. Specifically, individuals tended to select to be close to familiar nearest neighbours when moving slowly and, in general, selected to be closer to nearest neighbours in lichen habitat relative to forest and open habitats regardless of the familiarity of nearest neighbours. Within the movement ecology literature for ungulates, there is an assumption that slower movement in a given habitat represents foraging behaviour and faster movement represents searching behaviour (Owen-Smith et al., 2010). Our results support this assumption. Individuals moved more slowly in lichen habitat and moved more quickly in open habitats. Within a social context, individuals appear to collectively move through open habitat with familiar individuals, perhaps to new foraging patches. Individuals are more likely to trust social information about food sources and predation risk from familiar individuals, but the potential costs are an increase in competition at foraging patches. Individuals may balance the trade-off between competition and access to information by moving with socially familiar individuals but spacing apart during foraging. Lichen habitat is typically open, suggesting the possibility that individuals may remain in visual and vocal contact, thereby facilitating social cohesion during foraging despite physically spacing apart (Jacobs, 2010). This type of movement-dependent social association could contribute to the maintenance of social communities described above. Our results are also corroborated by other ungulate systems. In bison (*Bison bison*), the social environment in combination with recent knowledge of local foraging options dictated whether individuals followed, or left, a group (Merkle, Sigaud, &Fortin, 2015). Moreover, in the bison system, the costs and benefits of foraging in a group are moderated by collective decision making (Sigaud et al., 2017) and collective movement (Courant &Fortin, 2012), both of which are likely involved in the foraging decisions made by caribou. Here, we elucidate potential behavioural mechanisms (i.e., foraging or moving) that influence the frequency and magnitude of social associations.

The emergent geometry of collective movement and spatial arrangement of individuals in a group appears to change as individuals adjust their behaviour based on the availability of resources and the presence of familiar conspecifics (Morrell, Ruxton, &James, 2011). Assamese macaques (*Macaca assamensis*) distance from one another during foraging, but move collectively between foraging sites (Heesen, Macdonald, Ostner, &Schülke, 2015), while individual giraffes (*Giraffa camelopardalis*) show social preference for conspecifics during foraging, but not during movement (Muller et al., 2018). Interestingly, macaques foraged in closer proximity to individuals of similar dominance rank, but for giraffes it was unclear whether observed social preference was the result of passive or active assortment. For caribou, dominance hierarchies are linear and typically driven by body size (Barrette &Vandal, 1986), suggesting that social preference in caribou could also be related to dominance. Our ability to delineate aspects of the social environment between collective movement and habitat selection within a unified framework is useful for disentangling passive or active assortment, for example dominance rank, conspecific attraction, or the transfer of information about foraging resources.

We assumed that moving with familiar conspecifics is the result of information transfer about the location or quality of cratering sites, but spacing apart during foraging occurs because competition among individual caribou for craters in the winter can be substantial (Barrette &Vandal, 1986). Moreover, selection for open habitat relative to its availability in groups could also reflect the use of social information about the location of foraging sites (Lesmerises et al., 2018) or predation (Hamilton, 1971). Craters can vary in size and distribution (Bergerud, 1974); however, craters may only be large enough for a single individual to forage at a time (Mayor, Schaefer, Schneider, &Mahoney, 2009). Foraging apart from conspecifics reduce the costs of competition at cratering sites, which may be limited on the landscape or relatively small. We propose that while caribou generally have larger group sizes in winter (Webber &Vander Wal, 2021), groups vary in size based on movement and habitat selection behaviour presumably to balance the trade-off between competition and information acquisition. Furthermore, female caribou often have antlers, which unlike males, persist into winter. Females are hypothesized to use their antlers to defend craters and exert dominance over both males and females without antlers (Barrette &Vandal, 1986; Schaefer &Mahoney, 2001). This interpretation is corroborated by theory used to explain fission-fusion dynamics, where individuals are expected to split and merge through space and time to reduce conflict and competition during foraging.

We demonstrate assortment of individuals into distinct social communities, despite high range overlap with individuals in other communities. Integrating space and time revealed fine-scale processes that form social communities and the socially mediated nature of movement ecology and habitat selection. Within a unified socially informed integrated step selection framework, we bridge the theoretical and methodological gap between social network analysis, movement ecology, and habitat selection. We also demonstrate how social association is context-dependent, where individuals forage spaced apart from one another, but move collectively with familiar between foraging patches. Our synthesis of integrated step selection functions with social networks to test hypotheses is an important step towards identifying the roles of physical space and animal space use as factors influencing the social environment (Strandburg-Peshkin et al., 2017). Moreover, individual variation in phenotypes attributable to movement or habitat selection may affect how individuals experience the social environment (Webber et al., 2022; Webber &Vander Wal, 2018). Movement, habitat selection, and social behaviour are clearly linked; as van Moorter et al. (2016) described movement as the ‘glue’ connecting habitat selection to the physical location of a given set of habitats, we posit that movement is the glue connecting collective habitat selection to the social environment.

## ACKNOWLEDGEMENTS

We respectfully acknowledge the territory in which data were collected and analyzed as the ancestral homelands of the Beothuk and the Island of Newfoundland as the ancestral homelands of the Mi’kmaq and Beothuk. We thank M. Laforge, M. Bonar, J. Hendrix, and R. Huang for help in the field, and D. Wilson, K. Lewis, I. Fleming, G. Albery, members of the Wildlife Evolutionary Ecology Lab, and three anonymous reviewers for helpful comments on previous versions of this manuscript. We also thank members of the Newfoundland and Labrador Wildlife Division, including S. Moores, B. Adams, W. Barney, and J. Neville for facilitating animal captures and for logistical support in the field. Funding for this study was provided by Natural Sciences and Engineering Research Council (NSERC) Vanier Canada Graduate Scholarships to QMRW and CMP, a NSERC Canada Graduate Scholarship (Masters and Doctoral) to KAK, and a NSERC Discovery Grant to EVW.

## DATA AVAILABILITY STATEMENT

All code and data used for statistical analysis and figures are archived in Zenodo: doi:10.5281/zenodo.4549509

## AUTHORS CONTRIBUTIONS

QMRW and EVW conceived the ideas and designed method; QMRW conducted fieldwork QMRW and generated social networks; QMRW, CMP, KAK, and JWT conducted movement and spatial analysis; QMRW led the writing of the manuscript; EVW developed the research program. All authors contributed critically to drafts and gave final approval for publication.

## References

Avgar, T., Potts, J. R., Lewis, M. A., & Boyce, M. S. (2016). Integrated step selection analysis: Bridging the gap between resource selection and animal movement. Methods in Ecology and Evolution, 7(5), 619–630. doi: 10.1111/2041-210X.12528

Avgar, Tal, Lele, S. R., Keim, J. L., & Boyce, M. S. (2017). Relative Selection Strength: Quantifying effect size in selection inference. Ecology & Evolution, 7, 5322–5330. doi: 10.1002/ece3.3122

Barrette, C., & Vandal, D. (1986). Social rank, dominance, antler size, and access to food in snow-bound wild woodland caribou. Behaviour, 97(1), 118–145.

Basille, M., Fortin, D., Dussault, C., Bastille-Rousseau, G., Ouellet, J.-P., & Courtois, R. (2015). Plastic response of fearful prey to the spatio-temporal dynamics of predator distribution. Ecology, 96(10), 150511123704000. doi: 10.1890/14-1706.1

Bergerud, A. T. (1974). Relative abundance of food in winter for Newfoundland caribou. Oikos, 25(3), 379–387.

Bergerud, A. T., & Mercer, W. E. (1989). Caribou introductions in eastern North America. Wildlife Society Bulletin, 17(2), 111–120.

Best, E. C., Seddon, J. M., Dwyer, R. G., & Goldizen, A. W. (2013). Social preference influences female community structure in a population of wild eastern grey kangaroos. Animal Behaviour, 86(5), 1031–1040. doi: 10.1016/j.anbehav.2013.09.008

Bjørneraas, K., Van Moorter, B., Rolandsen, C. M., & Herfindal, I. (2010). Screening Global Positioning System Location Data for Errors Using Animal Movement Characteristics. Journal of Wildlife Management, 74(6), 1361–1366. doi: 10.2193/2009-405

Cairns, S. J., & Schwager, S. J. (1987). A comparison of association indices. Animal Behaviour, 35, 1454–1469.

Cantor, M., Wedekin, L. L., Guimarães, P. R., Daura-Jorge, F. G., Rossi-Santos, M. R., & Simões-Lopes, P. C. (2012). Disentangling social networks from spatiotemporal dynamics: The temporal structure of a dolphin society. Animal Behaviour, 84(3), 641–651. doi: 10.1016/j.anbehav.2012.06.019

Chamaillé-Jammes, S., Fritz, H., Valeix, M., Murindagomo, F., & Clobert, J. (2008). Resource variability, aggregation and direct density dependence in an open context: The local regulation of an African elephant population. Journal of Animal Ecology, 77(1), 135–144. doi: 10.1111/j.1365-2656.2007.01307.x

Courant, S., & Fortin, D. (2012). Time allocation of bison in meadow patches driven by potential energy gains and group size dynamics. Oikos, 121(7), 1163–1173. doi: 10.1111/j.1600-0706.2011.19994.x

Creel, S., Schuette, P., & Christianson, D. (2014). Effects of predation risk on group size, vigilance, and foraging behavior in an African ungulate community. Behavioral Ecology, 25(4), 773–784. doi: 10.1093/beheco/aru050

Croft, D. P., Darden, S. K., & Wey, T. W. (2016). Current directions in animal social networks. Current Opinion in Behavioral Sciences, 12, 52–58. doi: 10.1016/j.cobeha.2016.09.001

Csárdi, G., & Nepusz, T. (2006). The igraph software package for complex network research. InterJournal Complex Systems, 1695, 1–9.

Duchesne, T., Fortin, D., & Rivest, L. P. (2015). Equivalence between step selection functions and biased correlated random walks for statistical inference on animal movement. PLoS ONE, 10(4), 1–12. doi: 10.1371/journal.pone.0122947

Farine, D. R. (2013). Animal social network inference and permutations for ecologists in R using asnipe. Methods in Ecology and Evolution, 4(12), 1187–1194. doi: 10.1111/2041-210X.12121

Farine, D. R. (2014). Measuring phenotypic assortment in animal social networks: Weighted associations are more robust than binary edges. Animal Behaviour, 89, 141–153. doi: 10.1016/j.anbehav.2014.01.001

Farine, D. R., & Whitehead, H. (2015). Constructing, conducting and interpreting animal social network analysis. Journal of Animal Ecology, 84, 1144–1163. doi: 10.1111/1365-2656.12418

Fieberg, J., & Kochanny, C. O. (2005). Quanitfying home-range overlap: the importance of the utilization distribution. Journal of Wildlife Management, 69(4), 1346–1359. doi: 10.2193/0022-541X(2005)69

Fieberg, J., Rieger, R. H., Zicus, M. C., & Schildcrout, J. S. (2009). Regression modelling of correlated data in ecology: Subject-specific and population averaged response patterns. Journal of Applied Ecology, 46(5), 1018–1025. doi: 10.1111/j.1365-2664.2009.01692.x

Fortin, D., Beyer, H. L., Boyce, M. S., Smith, D. W., Duchesne, T., & Mao, J. S. (2005). Wolves influence elk movements: Behavior shapes a trophic cascade in Yellowstone National Park. Ecology, 86(5), 1320–1330.

Fortin, Daniel, Courtois, R., Etcheverry, P., Dussault, C., & Gingras, A. (2008). Winter selection of landscapes by woodland caribou: Behavioural response to geographical gradients in habitat attributes. Journal of Applied Ecology, 45(5), 1392–1400. doi: 10.1111/j.1365-2664.2008.01542.x

Giraldeau, L.-A., & Caraco, T. (2018). Social foraging theory (Vol. 73). Princeton University Press.

Giraldeau, L. A., & Dubois, F. (2008). Social foraging and the study of exploitative behavior. Advances in the Study of Behavior, 38(08), 59–104. doi: 10.1016/S0065-3454(08)00002-8

Hamilton, W. D. (1971). Geometry for the selfish herd. Journal of Theoretical Biology, 31(2), 295–311. doi: 10.1016/0022-5193(71)90189-5

Harel, R., Spiegel, O., Getz, W. M., & Nathan, R. (2017). Social foraging and individual consistency in following behaviour: testing the information centre hypothesis in free-ranging vultures. Proceedings of the Royal Society B, 284, 20162654.

He, P., Maldonado-Chaparro, A. A., & Farine, D. R. (2019). The role of habitat configuration in shaping social structure: a gap in studies of animal social complexity. Behavioral Ecology and Sociobiology, 73, 9.

Hebblewhite, M., & Merrill, E. (2008). Modelling wildlife-human relationships for social species with mixed-effects resource selection models. Journal of Applied Ecology, 45(3), 834–844. doi: 10.1111/j.1365-2664.2008.01466.x

Heesen, M., Macdonald, S., Ostner, J., & Schülke, O. (2015). Ecological and Social determinants of group cohesiveness and within-group spatial position in wild assamese macaques. Ethology, 121(3), 270–283. doi: 10.1111/eth.12336

Integrated-Informatics. (2014). Sustainable Development and Strategic Science Branch Land Cover Classification (Vol. 5, pp. 3–19). Vol. 5, pp. 3–19. St. John’s, NL.

Jacobs, A. (2010). Group cohesiveness during collective movements: Travelling apart together. Behavioural Processes, 84(3), 678–680. doi: 10.1016/j.beproc.2010.03.004

Jolles, J. W., King, A. J., & Killen, S. S. (2020). The role of individual heterogeneity in collective animal behaviour. Trends in Ecology & Evolution, 35, 278–291. doi: 10.1016/j.tree.2019.11.001

Jones, T. B., Patrick, S. C., Evans, J. C., & Wells, M. R. (2020). Consistent sociality but flexible social associations across temporal and spatial foraging contexts in a colonial breeder. Ecology Letters, 23, 1085–1096. doi: 10.1111/ele.13507

Kasozi, H., & Montgomery, R. A. (2020). Variability in the estimation of ungulate group sizes complicates ecological inference. Ecology and Evolution, 1–9. doi: 10.1002/ece3.6463

Lachlan, R. F., Crooks, L., & Laland, K. N. (1998). Who follows whom? Shoaling preferences and social learning of foraging information in guppies. Animal Behaviour, 56(1), 181–190. doi: 10.1006/anbe.1998.0760

Lesmerises, F., Johnson, C. J., & St-Laurent, M.-H. (2018). Landscape knowledge is an important driver of the fission dynamics of an alpine ungulate. Animal Behaviour, 140, 39–47. doi: 10.1016/j.anbehav.2018.03.014

Lima, S. L. (1995). Back to the basics of anti-predatory vigilance: the group-size effect. Animal Behaviour, 49, 11–20.

Mayor, S. J., Schaefer, J. A., Schneider, D. C., & Mahoney, S. P. (2009). The spatial structure of habitat selection: A caribou’s-eye-view. Acta Oecologica, 35(2), 253–260. doi: 10.1016/j.actao.2008.11.004

Merkle, J. A., Sigaud, M., & Fortin, D. (2015). To follow or not? How animals in fusion-fission societies handle conflicting information during group decision-making. Ecology Letters, 18(8), 799–806. doi: 10.1111/ele.12457

Morrell, L. J., Ruxton, G. D., & James, R. (2011). Spatial positioning in the selfish herd. Behavioral Ecology, 22(1), 16–22. doi: 10.1093/beheco/arq157

Mourier, J., Vercelloni, J., & Planes, S. (2012). Evidence of social communities in a spatially structured network of a free-ranging shark species. Animal Behaviour, 83(2), 389–401. doi: 10.1016/j.anbehav.2011.11.008

Muff, S., Signer, J., & Fieberg, J. (2020). Accounting for individual-specific variation in habitat-selection studies: Efficient estimation of mixed-effects models using Bayesian or frequentist computation. Journal of Animal Ecology, 89, 80–92. doi: 10.1111/1365-2656.13087

Muller, Z., Cantor, M., Cuthill, I. C., & Harris, S. (2018). Giraffe social preferences are context dependent. Animal Behaviour, 146, 37–49. doi: 10.1016/j.anbehav.2018.10.006

Newman, M. E. J. (2006). Modularity and community structure in networks. Proceedings of the National Academy of Sciences, 103(23), 8577–8582.

Owen-Smith, N., Fryxell, J. M., & Merrill, E. H. (2010). Foraging theory upscaled: the behavioural ecology of herbivore movement. Philosophical Transactions of the Royal Society B, 365(1550), 2267–2278. doi: 10.1098/rstb.2010.0095

Patin, R., Fortin, D., Sueur, C., & Chamaillé-Jammes, S. (2019). Space use and leadership modify dilution effects on optimal vigilance under food-safety trade-offs. The American Naturalist, 193(1), E15–E28. doi: 10.1086/700566

Peignier, M., Webber, Q. M. R., Koen, E. L., Laforge, M. P., Robitaille, A. L., & Vander Wal, E. (2019). Space use and social association in a gregarious ungulate: Testing the conspecific attraction and resource dispersion hypotheses. Ecology & Evolution, 9, 5133–5145. doi: 10.1002/ece3.5071

Pinter-Wollman, N., Hobson, E. A., Smith, J. E., Edelman, A. J., Shizuka, D., De Silva, S., … McDonald, D. B. (2013). The dynamics of animal social networks: Analytical, conceptual, and theoretical advances. Behavioral Ecology, 25(2), 242–255. doi: 10.1093/beheco/art047

Prokopenko, C. M., Boyce, M. S., & Avgar, T. (2017). Characterizing wildlife behavioural responses to roads using integrated step selection analysis. Journal of Applied Ecology, 54, 470–479. doi: 10.1111/1365-2664.12768

R Core Team. (2019). R: A language and environment for statistical computing.

Roberts, D. R., Bahn, V., Ciuti, S., Boyce, M. S., Elith, J., Guillera-Arroita, G., … Dormann, C. F. (2017). Cross-validation strategies for data with temporal, spatial, hierarchical, or phylogenetic structure. Ecography, 40(8), 913–929. doi: 10.1111/ecog.02881

Robitaille, A. L., Webber, Q. M. R., & Vander Wal, E. (2019). Conducting social network analysis with animal telemetry data: applications and methods using spatsoc. Methods in Ecology and Evolution, 10, 1203–1211. doi: 10.1111/2041-210X.13215

Schaefer, J. A., & Mahoney, S. P. (2001). Antlers on female caribou: Biogeography of the bones of contention. Ecology, 82(12), 3556–3560.

Shizuka, D., & Farine, D. R. (2016). Measuring the robustness of network community structure using assortativity. Animal Behaviour, 112, 237–246.

Shizuka, Daizaburo, Chaine, A. S., Anderson, J., Johnson, O., Laursen, I. M., & Lyon, B. E. (2014). Across-year social stability shapes network structure in wintering migrant sparrows. Ecology Letters, 17(8), 998–1007. doi: 10.1111/ele.12304

Sigaud, M., Merkle, J. A., Cherry, S. G., Fryxell, J. M., Berdahl, A., & Fortin, D. (2017). Collective decision-making promotes fitness loss in a fusion-fission society. Ecology Letters, 20(1), 33–40. doi: 10.1111/ele.12698

Signer, J., Fieberg, J., & Avgar, T. (2019). Animal movement tools (amt): R package for managing tracking data and conducting habitat selection analyses. Ecology and Evolution, 9(July 2018), 880–890. doi: 10.1002/ece3.4823

Silk, J. B. (2007). Social components of fitness in primate groups. Science, 317(5843), 1347–1351.

Spiegel, O., Leu, S. T., Bull, C. M., & Sih, A. (2017). What’s your move? Movement as a link between personality and spatial dynamics in animal populations. Ecology Letters, 20(1), 3–18. doi: 10.1111/ele.12708

Spiegel, O., Leu, S. T., Sih, A., & Bull, C. M. (2016). Socially interacting or indifferent neighbours? Randomization of movement paths to tease apart social preference and spatial constraints. Methods in Ecology and Evolution, 7, 971–979. doi: 10.1111/2041-210X.12553

Spiegel, Orr, & Crofoot, M. C. (2016). The feedback between where we go and what we know— information shapes movement, but movement also impacts information acquisition. Current Opinion in Behavioral Sciences, 12, 90–96. doi: 10.1016/j.cobeha.2016.09.009

Spiegel, Orr, Leu, S. T., Bull, C. M., & Sih, A. (2017). What’s your move? Movement as a link between personality and spatial dynamics in animal populations. Ecology Letters, 20(1), 3–18. doi: 10.1111/ele.12708

Spiegel, Orr, Sih, A., Leu, S. T., & Bull, C. M. (2018). Where should we meet? Mapping social network interactions of sleepy lizards shows sex-dependent social network structure. Animal Behaviour, 136, 207–215. doi: 10.1016/j.anbehav.2017.11.001

Strandburg-Peshkin, A., Farine, D. R., Crofoot, M. C., & Couzin, I. D. (2017). Habitat structure shapes individual decisions and emergent group structure in collectively moving wild baboons. ELIFE, 6, e19505. doi: 10.7554/eLife.19505

Strandburg-Peshkin, A., Papageorgiou, D., Crofoot, M. C., & Farine, D. R. (2018). Inferring influence and leadership in moving animal groups. Philosophical Transactions of the Royal Society B, 373, 20170006. doi: 10.1098/not

Sueur, C., King, A. J., Conradt, L., Kerth, G., Lusseau, D., Mettke-Hofmann, C., … Aureli, F. (2011). Collective decision-making and fission-fusion dynamics: A conceptual framework. Oikos, 120(11), 1608–1617. doi: 10.1111/j.1600-0706.2011.19685.x

van Beest, F. M., Uzal, A., Vander Wal, E., Laforge, M. P., Contasti, A. L., Colville, D., & Mcloughlin, P. D. (2014). Increasing density leads to generalization in both coarse-grained habitat selection and fine-grained resource selection in a large mammal. Journal of Animal Ecology, 83(1), 147–156. doi: 10.1111/1365-2656.12115

Van Moorter, B., Rolandsen, C. M., Basille, M., & Gaillard, J.-M. (2016). Movement is the glue connecting home ranges and habitat selection. Journal of Animal Ecology, 85, 21–31. doi: 10.1111/1365-2656.12394

Webber, Q.M.R., Albery, G. F., Farine, D. R., Pinter-Wollman, N., Sharma, N., Spiegel, O., … Manlove, K. (2022). Behavioural ecology at the spatial-social interface. EcoEvoRxiv, 1–40. Retrieved from 10.32942/osf.io/f7cm9

Webber, Q.M.R., & Vander Wal, E. (2018). An evolutionary framework outlining the integration of individual social and spatial ecology. Journal of Animal Ecology, 87(1), 113–127. doi: 10.1111/1365-2656.12773

Webber, Quinn M.R., Ferraro, K. M., Hendrix, J. G., & Vander Wal, E. (2022). What do caribou eat? A review of the literature on caribou diet. Canadian Journal of Zoology, 206(January), 197–206. doi: 10.1139/cjz-2021-0162

Webber, Quinn M R, & Vander Wal, E. (2021). Context-dependent group size: effects of population density, habitat, and season. Behavioral Ecology, 1–12. doi: 10.1093/beheco/arab070

Whitehead, H. (2008). Analyzing animal societies: quantitative methods for vertebrate social analysis. University of Chicago Press.

Worton, B. J. (1989). Kernel methods for estimating the utilization distribution in home-range studies. Ecology, 70(1), 164–168.

