## Supplementary materials for "Effects of the social environment on movement-integrated habitat selection"

**Appendix 1: additional information on study site and subjects**

(a) *Study site and subjects*

Newfoundland, as well as Fogo Island, has a humid-continental climate and persistent precipitation throughout the year. The dominant habitat types consisted of coniferous and mixed forests of balsam fir (*Abies balsamea*), black spruce (*Picea mariana*) and white birch (*Betula papyrifera*), as well as bogs, lakes, and barren rock. Fogo Island (237 km^2^) is situated ~12 km off the northeastern coast of Newfoundland (49º40’0’’ N, 54º11’0’’ W). Unlike many of the other herds in Newfoundland (Bastille-Rousseau, Schaefer, Mahoney, & Murray, 2013), caribou on Fogo Island have a relatively stable population that has not declined in recent years. In addition, caribou on Fogo Island are sedentary and do not display any migratory or long-distance movements. In winter, caribou activity budgets suggest that caribou spend ~50% of their time foraging, while ~40% of their time is spent lying down or ruminating, 7% of their time is spent walking or trotting, and 3% of their time is spent standing (Boertje, 1985; Duquette & Klein, 1987). Using Landsat images with 30m × 30m pixels (Integrated-Informatics, 2014), we determined the habitat composition of Fogo Island. Specifically, the Government of Newfoundland and Labrador developed a land classification system consisting of nine primary habitats, which we pooled for subsequent analyses (see main text). We pooled habitats into three categories. Open habitats that caribou move through consisted of wetland (21.2% of habitats on Fogo Island), rocky outcrops (6.7%), and water/ice (9.2%) habitats. Closed habitats consisted of conifer scrub (27.0%), conifer forest (10.4%), mixed-wood (9.2%), and broadleaf forest (0.2% of habitats) habitats. Finally, we considered lichen barrens (11.7%) as caribou foraging habitat. Overall, the proportion of GPS relocations (see below) that fell within each habitat were: 54.2% of GPS relocations were in open habitats, 30.4% of GPS relocations were in lichen habitat, 10.8% of GPS relocations were in forested habitats, and 4.4% of locations were in undesignated habitats.

We deployed GPS collars on 26 adult female caribou (n = 72 caribou-years) in three phases. In spring 2016, 2017 and 2018, collars were deployed on individual caribou (n = 13 in 2016, n = 11 in 2017, n = 1 in 2018). After two years, previously deployed collars were replaced, thus animals collared in spring 2016 were re-collared in spring 2018 (n = 11). Collars collected data throughout the year and were programmed to collect locations every two hours. For all analyses, we restricted locations to only include relocations from the first 75 days of each year (1 January–16 March). Prior to analyses, we removed all erroneous and outlier GPS locations following Bjørneraas et al. (2010). Specifically, we removed implausible GPS fixes, including those that were recorded in the ocean and those that exceeded the maximum diameter of Fogo Island (30 km). We also removed individuals with collar failure during the study period (n = 16 caribou-years) or individuals that swam to nearby adjacent islands (n = 3 caribou-years). After data screening, 24 adult female caribou (50 caribou-years) were used to generate annual social networks and 21 of these individuals (38 caribou-years) were used to assess patterns of movement, space use, and social behaviour in winter.

The sex ratio of caribou on Fogo Island is skewed because of male-biased hunting practices. We estimate that of the approximately 300 caribou that live on Fogo Island, there were ~180 adult females, ~50 adult males, and ~75 young of the year (Webber and Vander Wal *unpublished data*). In addition, all collared caribou in our study were adult females, indicating that the proportion of collared adult females was approximately 14.5% (26/180). For models of social network analysis, sensitivity analyses have identified that the greater the proportion of tagged animals from the whole population, the more robust model outcomes will be (Gilbertson, White, & Craft, 2021). However, in cases where tagging a high proportion of animals is logistically or financially impossible, other authors have called for an increase in the volume of data per individual included in the study (Silk, 2017). In the case of studies generated behavioural data using GPS collars, including ours, it is not possible to deploy collars on all individuals, and as a result, we rely on a relatively high volume of data per individual as opposed to a large proportion of tagged individuals. In our case, GPS collars were programmed to record the location of the individual every 2 hours (i.e., 12 locations per day × 75 days in our study = 900 potential relocations. In some cases, collars failed, or erroneous locations were recorded, and these locations were removed from subsequent analyses (see above). Overall, for the 38 caribou-years included in our social network analyses, we recorded an average of 886 observations per individual (SD = 28; range = 761–897). Thus, although the proportion of tagged adult females was only 14.5%, the volume of data per individual we used to generate social networks was relatively larger and consistently collected over a long period of time (i.e., relocations every 2 hours over 75 days).

**Appendix 2: Habitat selection analysis validation**

(a) *Model validation: k-fold cross validation*

We evaluated the predictive performance of our models using an out of sample discrimination k-fold cross validation test (Boyce, Vernier, Nielsen, & Schmiegelow, 2002; Roberts et al., 2017). We followed methods for conditional logistic regression models presented by Fortin et al., (2009). K-fold cross validation is a procedure which While resource selection functions that do not use conditional logistic regression subset *all* data, to generate k-folds, models fit with a conditional logistic regression require strata to be subset, as opposed to all data. Specifically, using a random sample of 80% of the available strata (i.e. one observed and 20 control steps), we built a new iSSF model including the same fixed and random effect variables as our original model and used the resulting parameter estimates to predict iSSF scores for the observed and control steps in the remaining 20% of the strata. We performed five folds, withholding a different 20% of strata for testing each time. Based predicted iSSF scores, we ranked observed and control steps from 1 to 21 and tallied the ranks of observed steps into 21 potential bins. We then performed Spearman rank correlation between the bin’s ranking and its associated frequency. Positive correlations therefore indicate increasing relative use in the test sample as a function of predicted values from the model fit using the training data. The mean and SD are presented in the main text.

**Table S1.** Summary of winter community assignment for three caribou social networks (2017–2019), including the number of individuals (N), number of communities, community size, modularity (*Q*), and the community assortativity coefficient (R_com_ = 0 indicates no confidence in community assignment; R_com_ = 1 indicates certainty in community assignment).

| **Year** | **N** | **Community** | **Community size** | **Home range area (km^2^)** | ***Q*** | **R_com_** |
| --- | --- | --- | --- | --- | --- | --- |
| 2017 | 10 | 1 | 3 | 84.71 | 0.13 | 0.95 |
|  |  | 2 | 7 | 43.43 |  |  |
| 2018 | 16 | 1 | 5 | 113.09 | 0.17 | 1.00 |
|  |  | 2 | 7 | 77.31 |  |  |
|  |  | 3 | 1 | 13.84 |  |  |
|  |  | 4 | 1 | 23.11 |  |  |
|  |  | 5 | 1 | 4.94 |  |  |
|  |  | 6 | 1 | 19.06 |  |  |
| 2019 | 12 | 1 | 2 | 32.39 | 0.13 | 1.00 |
|  |  | 2 | 9 | 33.87 |  |  |
|  |  | 3 | 1 | 1.26 |  |  |

**Table S2.** Summary of Utilization Distribution Overlap Index (UDOI) for caribou social community across three years. Note, although some communities did not overlap, these were generally restricted to communities that contained a single individual (communities 5, 6, 7, and 8 in 2018; see Table 2 of the main text for details).

| **Comparison** | **Year** | **UDOI** |
| --- | --- | --- |
| Communities 1–2 | 2017 | 0.689 |
| **Annual average** | **2017** | **0.689** |
| Communities 1–2 | 2018 | 0.857 |
| Communities 1–3 | 2018 | 0.357 |
| Communities 2–3 | 2018 | 0.373 |
| Communities 1–4 | 2018 | 0.445 |
| Communities 2–4 | 2018 | 0.482 |
| Communities 3–4 | 2018 | 0.830 |
| Communities 1–5 | 2018 | 0 |
| Communities 2–5 | 2018 | 0 |
| Communities 3–5 | 2018 | 0 |
| Communities 4–5 | 2018 | 0 |
| Communities 1–6 | 2018 | 0.408 |
| Communities 2–6 | 2018 | 0.436 |
| Communities 3–6 | 2018 | 0.844 |
| Communities 4–6 | 2018 | 0.980 |
| Communities 5–6 | 2018 | 0 |
| **Annual average** | **2018** | **0.40 ± 0.35** |
| Communities 1–2 | 2019 | 0.286 |
| Communities 1–3 | 2019 | 0 |
| Communities 2–3 | 2019 | 0 |
| **Annual average** | **2019** | **0.095 ± 0.16** |
| **Total average** | **All years** | **0.37 ± 0.34** |

**Table S3.** Summary of top rated integrated step selection function based on model selection ($\Delta$AICc = 70969). For covariates with 95% confidence intervals that do not overlap zero we provide a brief interpretation of the result.

| Covariate | ß (95% CI) | z-value | p-value | Interpretation |
| --- | --- | --- | --- | --- |
| Intercept | –4.36 (–15.2, 6.45) | –0.79 | 0.43 | – |
| Step length | 1.42 (–0.16, 0.19) | 0.18 | 0.89 | – |
| Proportion Forest | 12.0 (10.2, 13.9) | 12.7 | <0.0001 | Individuals select areas with a higher proportion of forest habitat relative to the availability of forest habitat. |
| Proportion Lichen | 4.49 (2.67, 6.32) | 4.83 | <0.0001 | Individuals select areas with a higher proportion of lichen habitat relative to the availability of lichen habitat. |
| Proportion Open | 11.9 10.3, 13.5) | 14.7 | <0.0001 | Individuals select areas with a higher proportion of open habitat relative to the availability of open habitat. |
| Nearest neighbour distance (end) | –0.24 (–0.36, –0.11) | –3.81 | 0.0001 | Individuals end their steps close to nearest neighbours. |
| Simple ratio index | –1.01 (–1.95, –0.08) | 2.12 | 0.03 | Nearest neighbours at the end of a step have lower shared values of the SRI. |
| Proportion Forest : Step length | –0.07 (–0.17, 0.02) | –1.48 | 0.13 | – |
| Proportion Lichen : Step length | –0.12 (–0.22, –0.03) | –2.52 | 0.01 | Individuals take shorter steps when selecting lichen habitat relative to the availability of lichen habitat. |
| Proportion Open : Step length | 0.32 (0.25, 0.41) | 8.03 | <0.0001 | Individuals take longer steps when selecting open habitat relative to the availability of open habitat. |
| Step length : Nearest neighbour distance (start) | –0.012 (–0.04, 0.01) | –0.95 | 0.34 | – |
| Proportion Forest : Nearest neighbour distance (end) | –0.49 (–0.63, –0.35) | –6.84 | <0.0001 | Individuals are closer to their nearest neighbour when selecting forest habitat relative to the availability of forest habitat. |
| Proportion Lichen : Nearest neighbour distance (end) | 0.07 (–0.05, 0.20) | 1.14 | 0.26 | – |
| Proportion Open : Nearest neighbour distance (end) | –0.55 (–0.67, –0.44) | –9.31 | <0.0001 | Individuals are closer to their nearest neighbour when selecting open habitat relative to the availability of open habitat. |
| Step length : Simple ratio index | –0.05 (–0.11, 0.01) | –1.64 | 0.10 | – |
| Proportion Forest : Simple ratio index | 5.55 (4.53, 6.56) | 10.7 | <0.0001 | Individuals share a higher dyadic SRI value with their nearest neighbour when selecting forest habitat relative to the availability of forest habitat. |
| Proportion Lichen : Simple ratio index | 1.47 (0.47, 2.47) | 2.88 | 0.0004 | Individuals share a higher dyadic SRI value with their nearest neighbour when selecting lichen habitat relative to the availability of lichen habitat. |
| Proportion Open : Simple ratio index | 6.02 (5.15, 6.90) | 13.5 | <0.0001 | Individuals share a higher dyadic SRI value with their nearest neighbour when selecting open habitat relative to the availability of open habitat. |

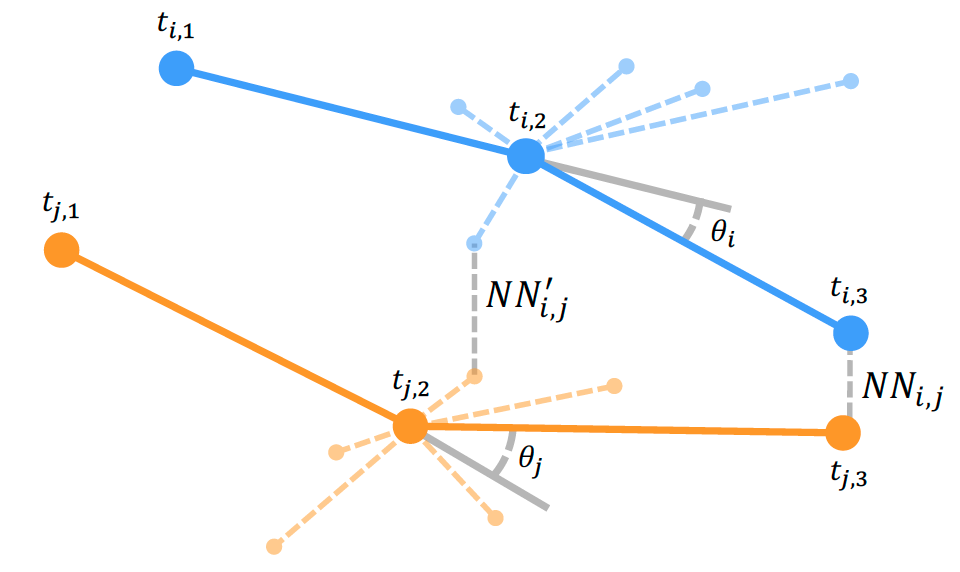

**Figure S1.** Schematic of integrated step selection functions in the context of social network analysis. Available (random) steps are generated based on the distribution of used step lengths (thin dashed orange and blue lines) and turn angles. We compared used (observed) to available (random) nearest neighbour distance based on fine-scale movement decisions of individuals. Blue lines represent used (dark thick lines) and available (light dashed lines) steps of individual *i* and orange lines represent used (dark thick lines) and available (light dashed lines) steps of individual *j.* The dashed grey line $({NN}_{i,j}$) represents the observed nearest neighbour distance between *i* and *j* at *t_3_*. For each set of available steps, we re-calculated nearest neighbour distance, denoted by a dashed grey line (${NN}_{i,j}^{'}$) between available steps for *i* and *j*, which represents the available nearest neighbour distance at a given iteration. Step length is the distance between the used start (e.g. *t_j,2_*) point and the step end point (*t_j,3_*) and turn angle is calculated as the angular deviation between the previous step heading (grey line and $\theta_{i}$ and $\theta_{j}$) and the subsequent used step.

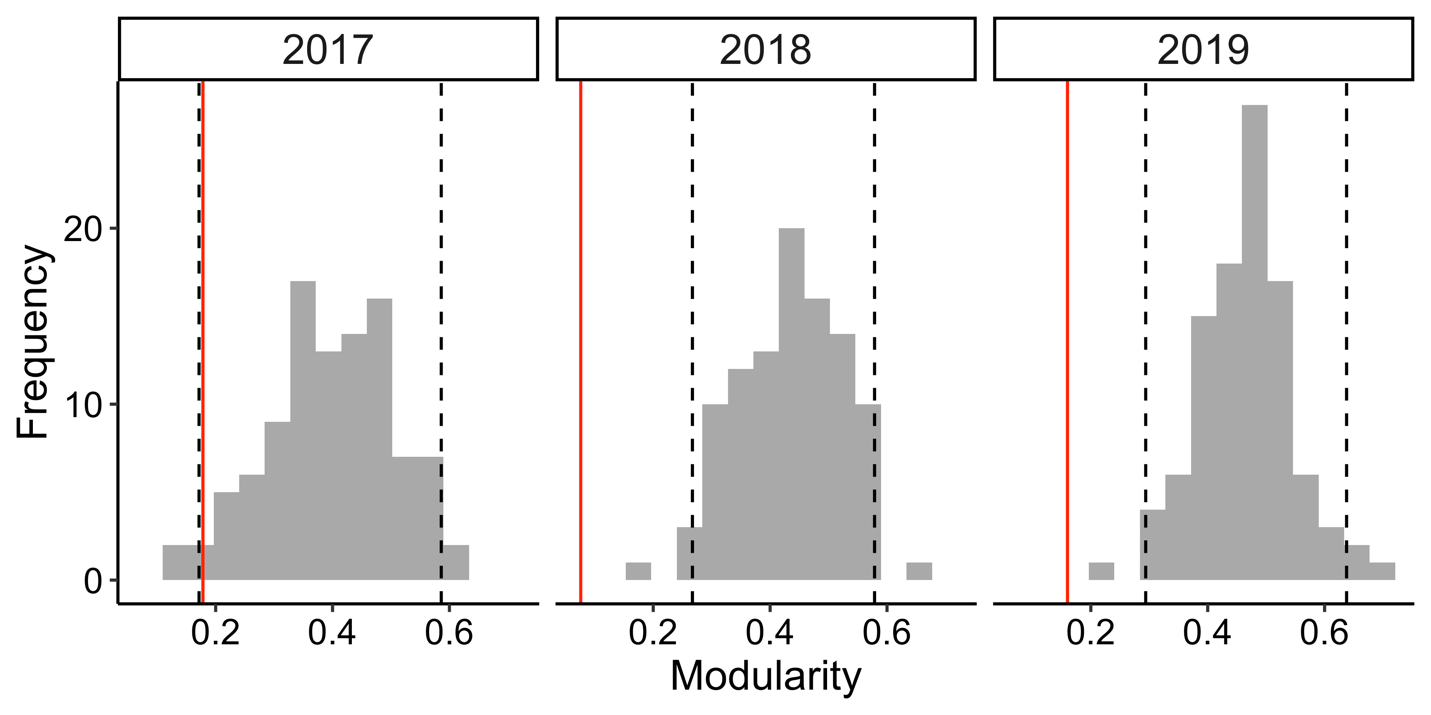

**Figure S2.** Comparison of observed modularity values (vertical red line) to the distribution of modularity values generated from null models in each year (95% confidence intervals are represented in each year by dashed black lines). In all years, observed modularity values were lower than the null distribution suggesting social associations among individuals in different social communities were more likely than expected by chance.

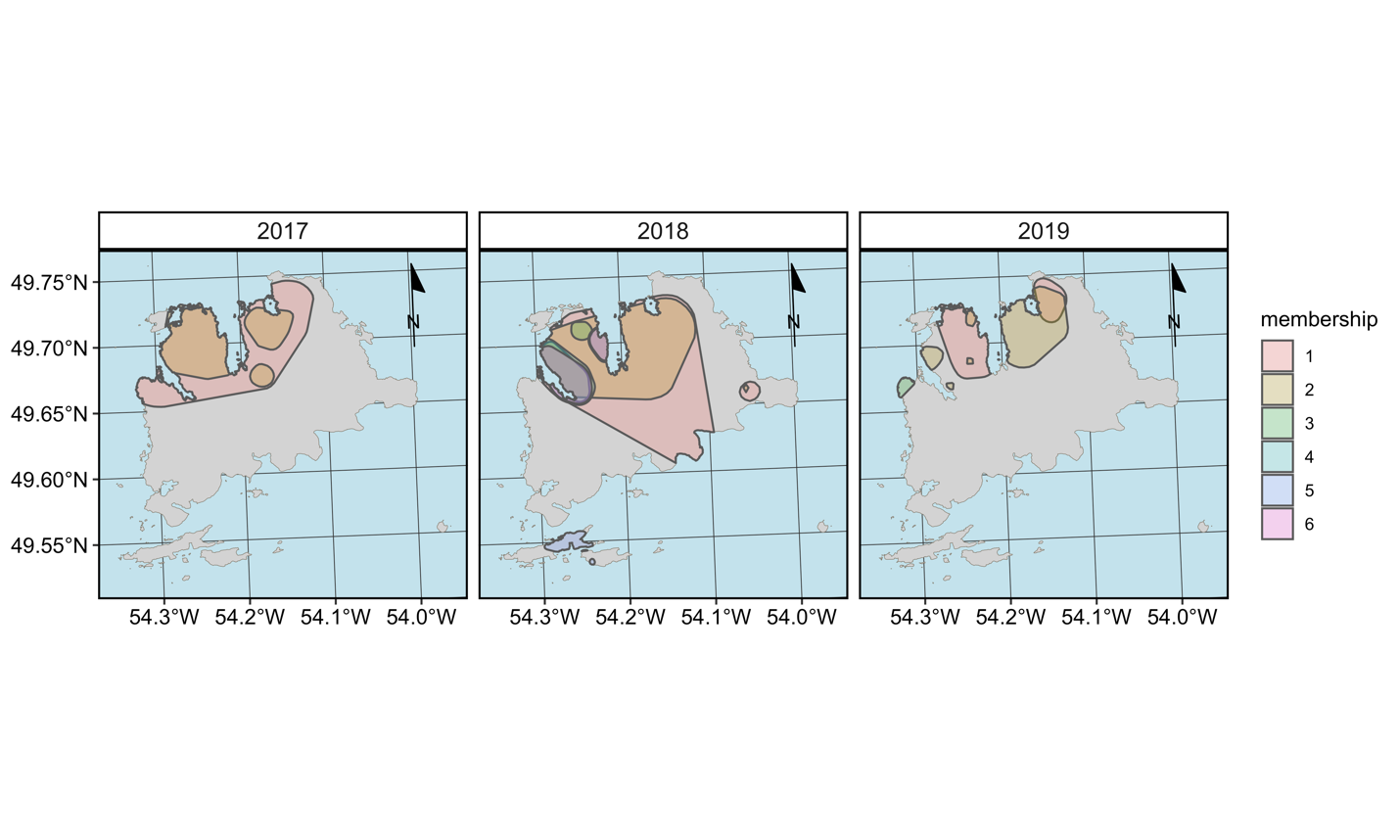

**Figure S3.** 95% kernel density estimates for Fogo Island social communities (2017–2019). Note, community overlap was relatively high in all years (Table S3).

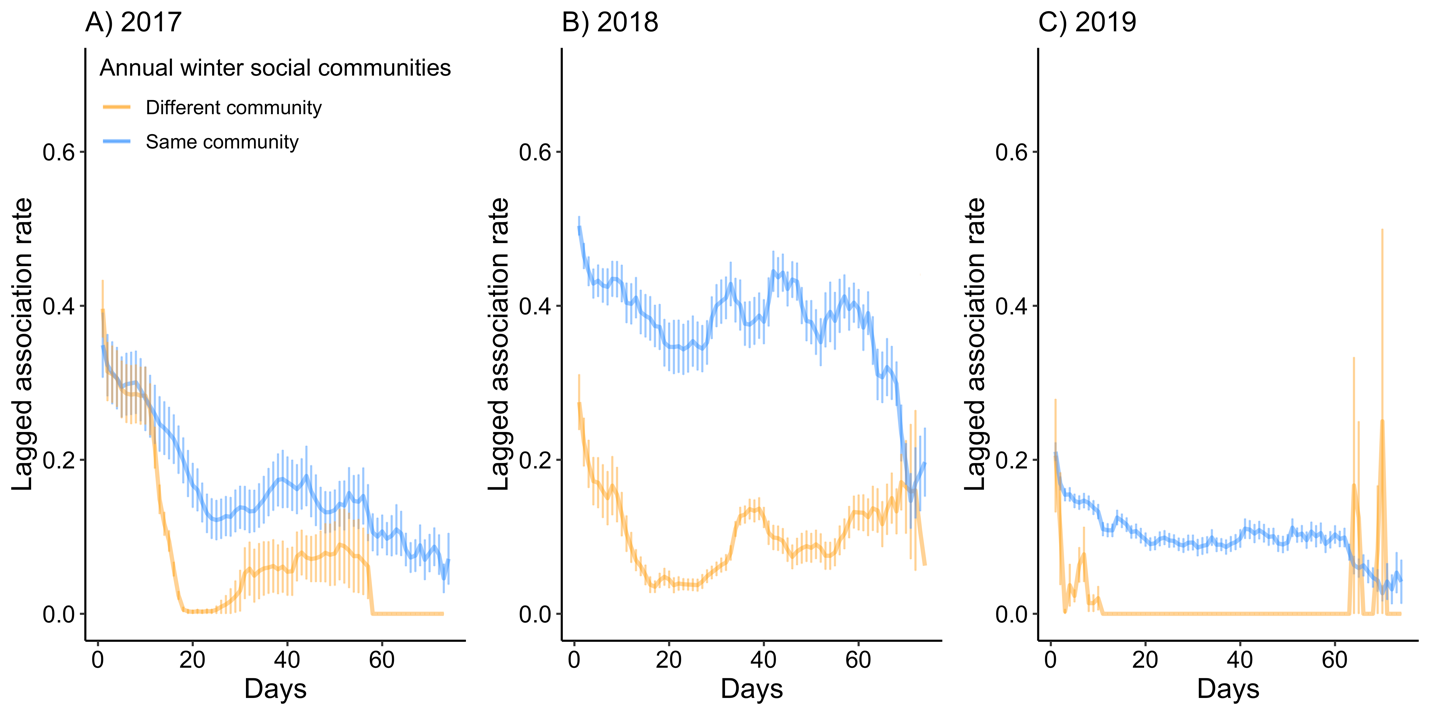

**Figure S4.** Observed lagged association rate (LAR) for caribou in the same (blue lines) and different (orange) annual winter social communities, calculated as the probability that any pair of individuals associated on a given day, are still associated on subsequent days. Note, the time period for LAR analysis and social community assignment was 1 January to 16 March. Error bars represent the standard error of all pairwise association rates calculated on each day. Individuals in the same social communities (blue lines) generally had higher lagged association rates, suggesting they were more likely to associate together over time.

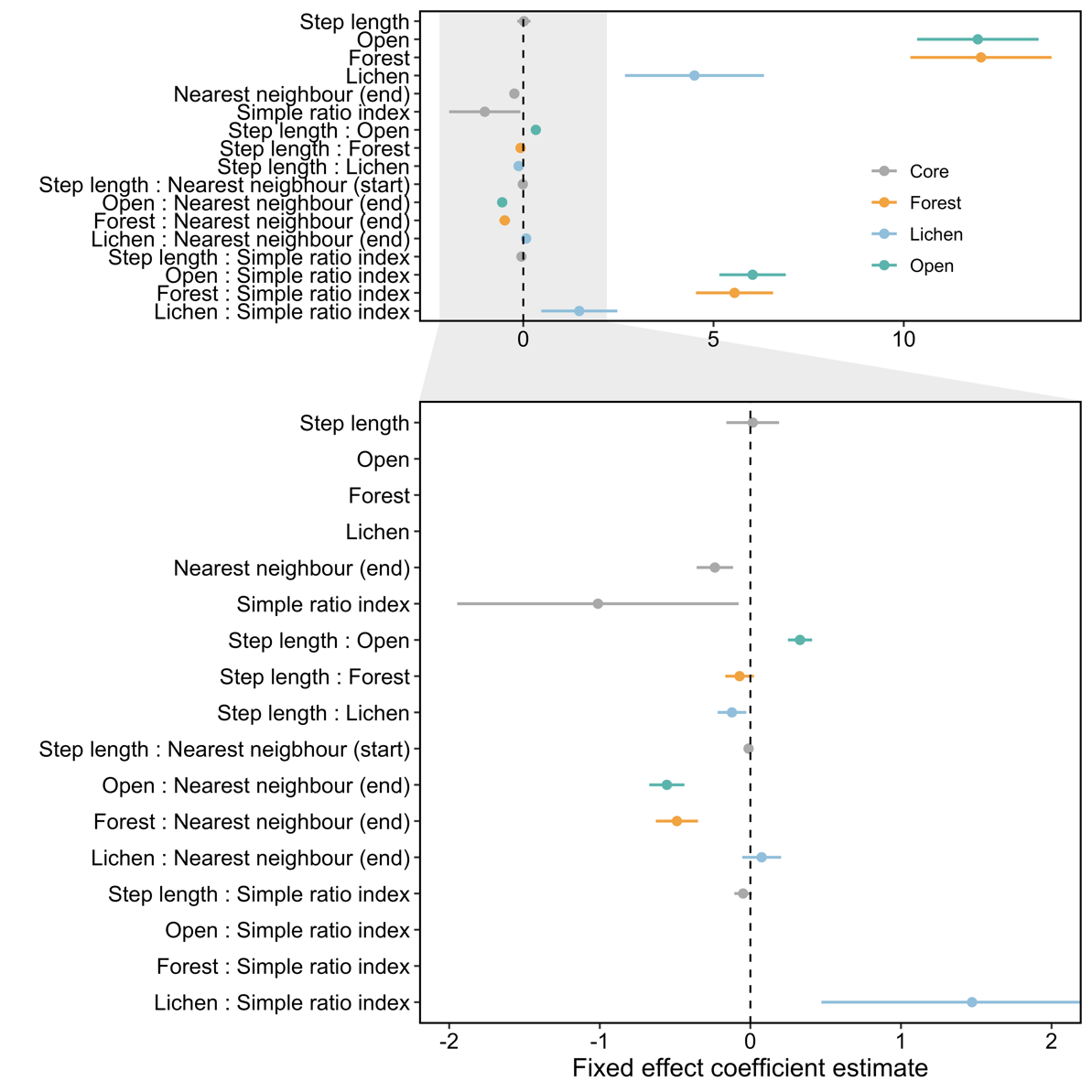

**Figure S5.** Habitat selection coefficients and 95% confidence intervals for our integrated step selection model. Coefficients that do not overlap zero are deemed significant (see Table S3). Colours denote which habitat each fixed effect term was associated with and in cases where a term was not associated with habitat, points were coloured grey. Step length, nearest neighbour distances (both start and end), and the simple ratio index were log-transformed. Measures of distance at the end of a step are included when the distance term is a single term in a model or included in an interaction with habitat type, but measures of distance at the end of the step are included when the distance term is in an interaction with step length. These terms model the effect of nearest neighbour distance at the end of a step on the likelihood of selecting a given habitat type because in the case of habitat selection, proximity to conspecifics is more relevant for where an animal is going. By contrast, the effect of nearest neighbour distance at the start of a step on the likelihood of taking a shorter or longer step because in the case of movement, proximity to conspecifics is more relevant for where an animal is starting. Note, the lower panel is a zoom of the upper panel for enhanced visualization of coefficients with relatively small confidence intervals.

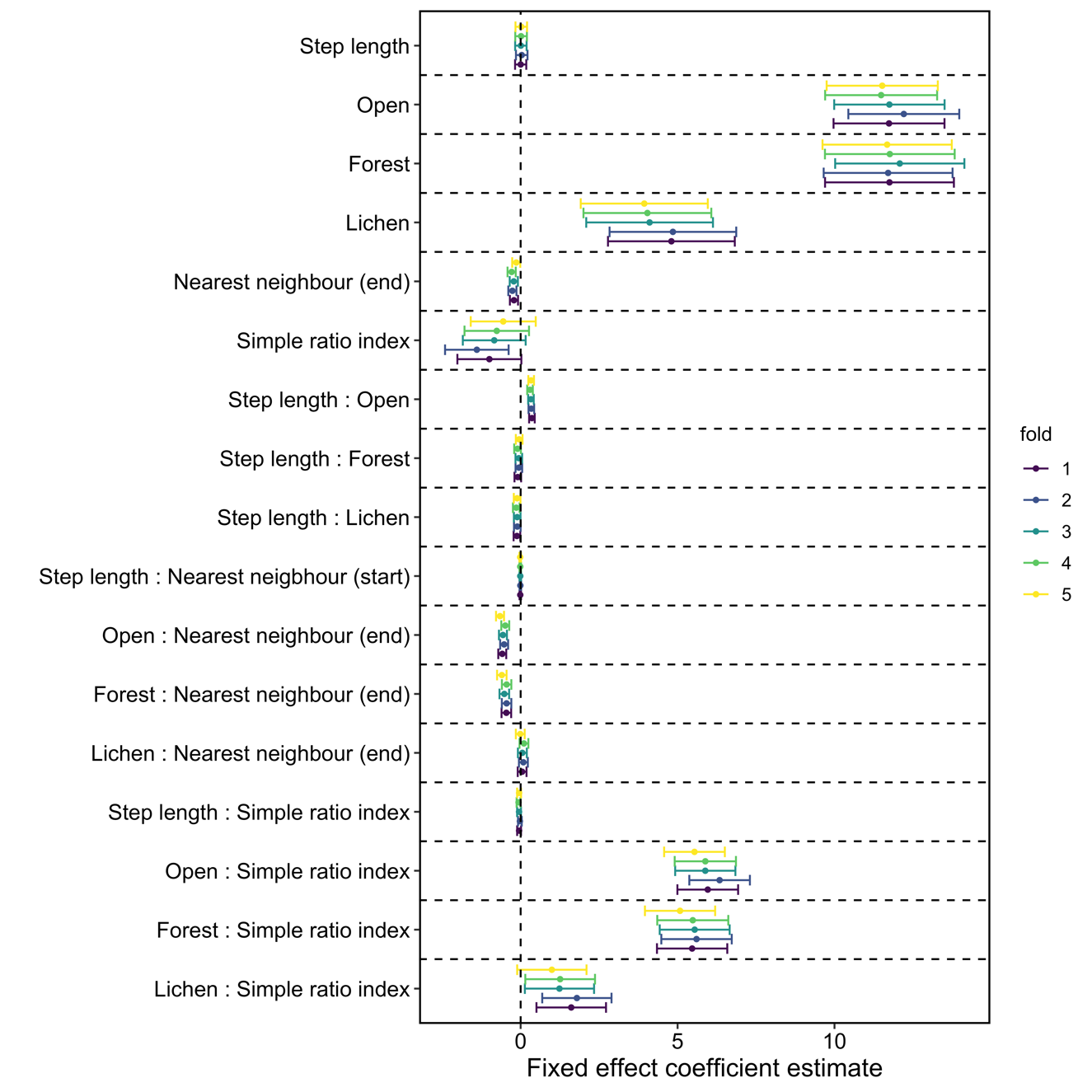

**Figure S6:** Summary of integrated step selection function coefficients from k-fold cross validation. Coefficient confidence intervals for terms in each k-fold overlap, therefore suggesting identical patterns of interpretation for each fold. We are therefore confident our model results are robust.

**References**

Bastille-Rousseau, G., Schaefer, J. A., Mahoney, S. P., & Murray, D. L. (2013). Population decline in semi-migratory caribou (Rangifer tarandus): intrinsic or extrinsic drivers? *Canadian Journal of Zoology*, *91*(11), 820–828. doi: doi:10.1139/cjz-2013-0154

Bjørneraas, K., Van Moorter, B., Rolandsen, C. M., & Herfindal, I. (2010). Screening global positioning system location data for errors using animal movement characteristics. *Journal of Wildlife Management*, *74*(6), 1361–1366. doi: 10.2193/2009-405

Boertje, R. D. (1985). An Energy Model for Adult Female Caribou of the Denali Herd, Alaska. *Journal of Range Management*, *38*(5), 468–473. doi: 10.2307/3899725

Boyce, M. S., Vernier, P. R., Nielsen, S. E., & Schmiegelow, F. K. . (2002). Evaluating resource selection functions. *Ecological Modelling*, *157*(2–3), 281–300. doi: 10.1016/S0304-3800(02)00200-4

Duquette, L. S., & Klein, D. R. (1987). Activity budgets and group size of caribou during spring migration. *Canadian Jounal of Zoology*, *65*(164–168).

Fortin, D., Fortin, M. E., Beyer, H. L., Duchesne, T., Courant, S., & Dancose, K. (2009). Group-size-mediated habitat selection and group fission-fusion dynamics of bison under predation risk. *Ecology*, *90*(9), 2480–2490. doi: 10.1890/08-0345.1

Gilbertson, M. L. J., White, L. A., & Craft, M. E. (2021). Trade-offs with telemetry-derived contact networks for infectious disease studies in wildlife. *Methods in Ecology and Evolution*, *12*(1), 76–87. doi: 10.1111/2041-210X.13355

Integrated-Informatics. (2014). *Sustainable Development and Strategic Science Branch Land Cover Classification* (Vol. 5, pp. 3–19). Vol. 5, pp. 3–19. St. John’s, NL.

Roberts, D. R., Bahn, V., Ciuti, S., Boyce, M. S., Elith, J., Guillera-Arroita, G., … Dormann, C. F. (2017). Cross-validation strategies for data with temporal, spatial, hierarchical, or phylogenetic structure. *Ecography*, *40*(8), 913–929. doi: 10.1111/ecog.02881

Silk, M. J. (2017). The next steps in the study of missing individuals in networks: a comment on Smith et al. (2017). *Social Networks*. doi: 10.1016/j.socnet.2017.05.002
